# Flapping at Resonance: Measuring the Frequency Response of the Hymenoptera Thorax

**DOI:** 10.1101/2019.12.11.873562

**Authors:** Mark A. Jankauski

## Abstract

Insects with asynchronous flight muscles are believed to flap at the fundamental frequency of their thorax or thorax-wing system. Flapping in this manner leverages the natural elasticity of the thorax to reduce the energetic requirements of flight. However, to the best of our knowledge, the fundamental frequency of the insect thorax has not been measured via vibration testing. Here, we measure the linear frequency response function (FRF) of several Hymenoptera (*Apis mellifera, Polistes dominula, Bombus huntii*) thoraxes about their equilibrium states in order to determine their fundamental frequencies. FRFs relate the input force to output acceleration at the insect tergum and are acquired via a mechanical vibration shaker assembly. When compressed 50 *μm*, thorax fundamental frequencies in all specimens approximately 50-150% higher than reported wingbeat frequencies. We suspect that the measured fundamental frequencies are higher in the experiment than during flight due to experimental boundary conditions that stiffen the thorax. Thus, our results corroborate the idea that some insects flap at the fundamental frequency of their thorax. Next, we compress the thorax between 100 - 300 μm in 50 μm intervals to assess the sensitivity of the fundamental frequency to geometric modifications. For all insects considered, the thorax fundamental frequency increased nearly monotonically with respect to level of compression. This implies that the thorax behaves a nonlinear hardening spring, which we confirmed via static force-displacement testing. Hardening behavior may provide a simple mechanism for the insect to adjust wingbeat frequency, and implies the thorax may behave as a nonlinear Duffing oscillator excited at large amplitude. The Duffing oscillator exhibits amplitude-dependent resonance and may serve as a useful model to increase the flapping frequency bandwidth of small resonant-type flapping wing micro air vehicles.

## 1. Introduction

Insects are remarkable fliers. They are capable of hovering [1], performing aggressive aerial maneuvers [2] and landing on surfaces in various orientations [3]. Given these capabilities, flying insects often serve as inspiration for miniature flapping wing micro air vehicles (FWMAVs) [4–6]. Insects have guided the design of FWMAV wings [7], actuators [8] and sensing modalities [9,10]. While early designs have shown potential, FWMAVs continue to face technical challenges that must be overcome before they are suitable for widespread implementation. Among these challenges is improving vehicle energy efficiency to enable autonomous flight. Presently, many FWMAVs are tethered and require off-board sources to provide power [11]. A comprehensive understanding of the insect “drivetrain”, in particular the insect thorax, may guide the design of energy efficient FWMAVs moving forward.

The thorax is the central component of the insect flight drivetrain (Fig. 1). It is an approximately ellipsoidal structure that contains two large sets of muscles called the dorsoventral (DVMs) and dorsolongitudinal muscles (DLMs) [12]. As these muscles contract, they deform the thorax which indirectly causes the insect’s wings to flap via an elaborate linkage mechanism called the wing hinge [13]. The insect may engage fine steering muscles in the hinge to maneuver [12]. Many insects leverage indirect wing actuation to flap [14], which in some cases is believed to reduce the power requirements of flight [15].

**Figure 1:**
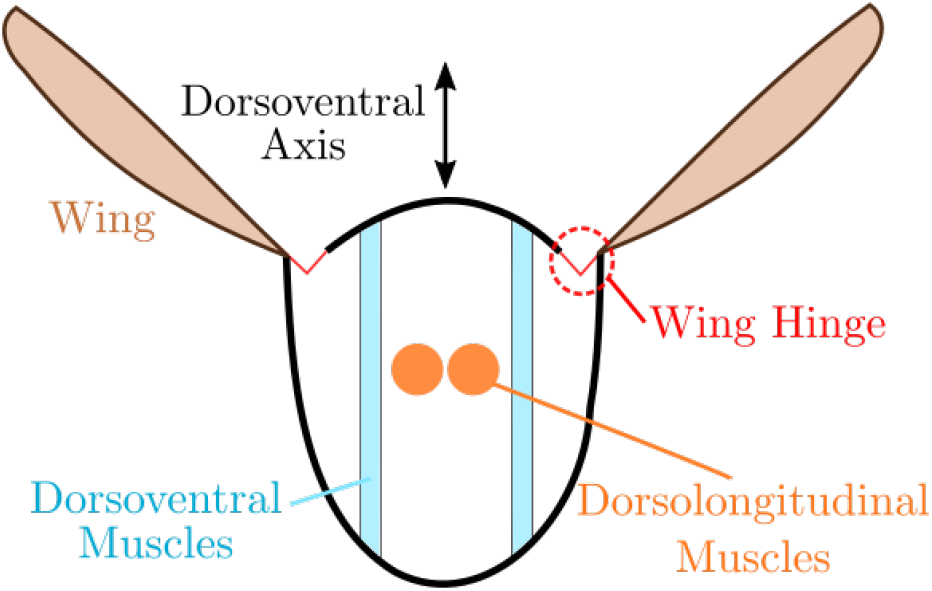
Simplified schematic of an insect thorax cross section.

In addition to indirect actuation, many insects utilize a unique type of “asynchronous” flight muscle to reduce the energetic costs of locomotion [16]. Unlike synchronous muscles, there is no one-to-one ratio between between muscle stimulation and mechanical contraction in asynchronous muscles – instead, they fire only once to initiate several wing beats. The dynamic properties of the thorax, rather than periodic muscle signaling, cause the muscle to repeatably contract in a process called “stretch activation” [14]. Interestingly, this muscle type has evolved independently numerous times along different branches of the insect phylogenetic tree, and researchers estimate the DVM and DLM flight muscles are asynchronous in approximately 75% of flying insects [16].

Much of the seminal research on asynchronous muscles was conducted by JWS Pringle in the 1950’s and 1960’s [17–20]. Pringle first observed that flight muscle length oscillations were higher and independent in frequency compared to neuron firing rate in the blowfly *Calliphora erythrocephala* [14]. Later, Pringle and colleagues attached a cyber-physical system with simulated mass, stiffness and damping to the asynchronous flight muscle of the Indian rhinoceros beetle *Oryctes rhinoceros* [18]. They found that the insect’s muscle contraction frequency was equivalent to the fundamental frequency of the cyber-physical system to which it was attached. This suggested that the insect thorax, or thorax-wing assembly, must have a resonant frequency equivalent to the insect’s wingbeat frequency to support asynchronous muscle function. However, some insects vary their wingbeat frequency during flight [21,22], whereas linear mechanical systems typically have fixed resonant frequencies. To explain this, researchers suggested that steering muscles could be used to stiffen the thorax [23, 24]. Stiffening would increase the resonant frequency of the thorax, and as a result the insect’s wingbeat frequency.

### 1.1. Scope of Present Research

Despite indirect evidence that insect’s flap at the fundamental frequency of their thorax, to the best of our knowledge the frequency response function (FRF) of the thorax has never been measured. Frequency response analysis, in the mechanical sense, involves exciting a structure over a wide frequency range with a known input signal and measuring the structure’s response. The ratio between input and output in the frequency domain is the frequency response function (FRF). Spectral peaks in the FRF typically correspond to the structure’s natural frequencies.

Given the motivation, the goals of the present work are to (1) determine the thorax FRF and to identify its fundamental frequency, and (2) assess if geometric modifications to the thorax achieved via compression cause its natural frequencies to change. The first objective will indicate if some insects indeed flap at the fundamental frequency of their thorax. To be clear, we refer to the first natural frequency of the thorax as its fundamental frequency. The second objective will probe the sensitivity of the thorax’s natural frequency to thorax shape, which could describe a potential mechanism for an insect to change wingbeat frequency. This work is restricted to describe only the small amplitude *linear* behavior of the thorax about varying equilibrium states. Nonlinear behavior is not considered.

The remainder of this paper is organized as follows. We first detail the custom experimental setup used for dynamic testing as well as insect specimen collection and preparation. For all experimental studies we use insects from the order Hymenoptera, all of which have asynchronous flight muscles. Specifically, we use the honeybee *Apis mellifera*, European paper wasp *Polistes dominula*, and bumblebee *Bombus huntii*. We then analyze FRF data and discuss the dynamic behavior of the thorax. Based upon these findings, we conduct a simple static analysis on the honeybee thorax. We conclude by examining the implications of this research.

## 2. Methods

We constructed a custom shaker assembly that allows us to excite an insect thorax about its dorsoventral axis (Fig. 2). Using this assembly, we determine the FRF *G*(*jω*) that relates the force *F*(*jω*) applied at a point on the ventral surface of the thorax to the acceleration *A*(*jω*) at the same point, where *ω* denotes excitation frequency and j denotes the imaginary unit. We define force as the input and acceleration as the output such that 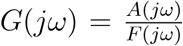. A mechanical idealization of the system is shown in Fig. 3. We are able to determine the thorax’s fundamental frequency via peaks in the FRF magnitude.

**Figure 2:**
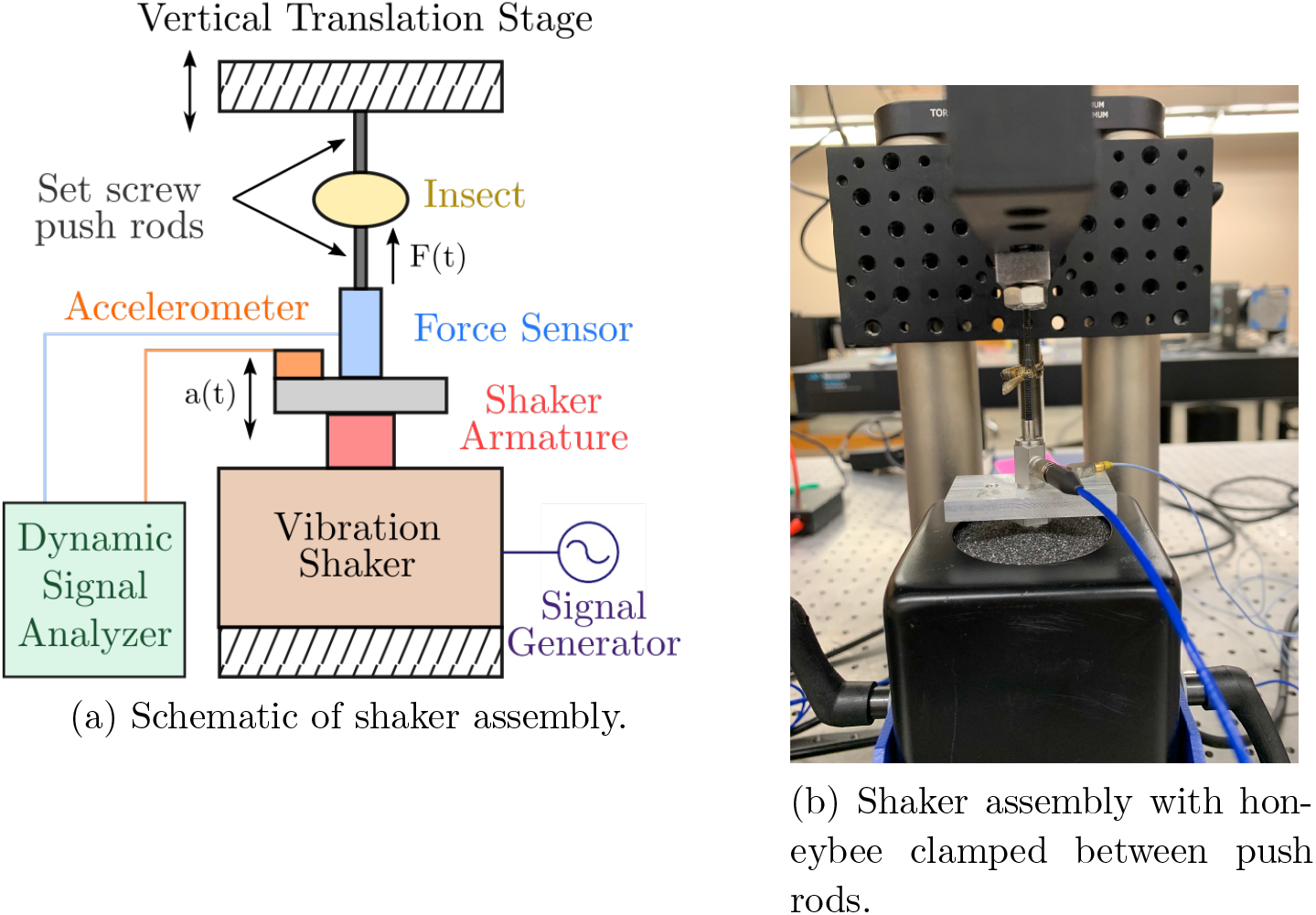
Experimental assembly used for measuring the insect thorax FRF.

**Figure 3:**
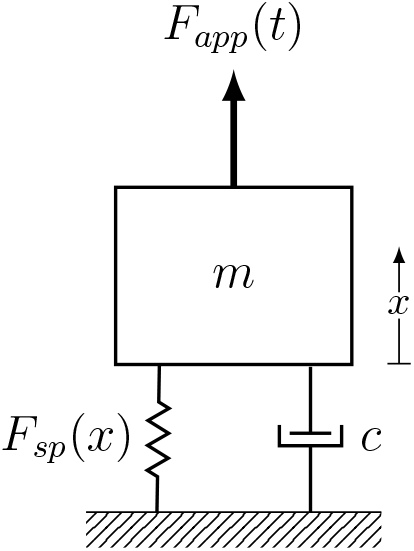
Mechanical representation of FRF experiment. *F_app_* denotes the force to the shaker, *x* denotes thorax displacement, *F_sp_*(*x*) denotes the thorax restorative force and *c* denotes a damping coefficient.

### 2.1. Dynamic Experimental Setup

We use an electrodynamic vibration shaker (The Modal Shop, 2007E) rated at 31 N and powered by an external amplifier (The Modal Shop, 2100E21-100) to excite the thorax. A high-sensitivity piezoelectric force sensor (PCB Piezotronics, 209C11) threads directly into the shaker’s insert. Then, a piezoelectric accelerometer (PCB Piezotronics, 352A21) is mounted via wax to a thick plate between the force sensor and shaker. The force sensor and accelerometer measure the input and output respectively, and are both powered by a benchtop signal conditioner (PCB Piezotronics, 482C15).

We then place a large, coarse vertical translation stage (Thorlabs, VAP10) with a vertically mounted optical breadboard behind the shaker assembly (Fig. 2b). To enable fine vertical adjustment, we mount an additional one-axis translation stage (Newport, UMR8.25 stage with BM17.25 micrometer, 1.0 *μm* resolution) to the vertical breadboard. An L-bracket is cantilevered off the fine linear stage. We thread a small push rod (#2 screws) into a metal mounting plate, and attach this mounting plate directly to the L-bracket. We then thread a second #2 screw directly into the force cell that is attached to the shaker. The two screws are colinear such that when the vertical stage is lowered, the insect thorax is compressed between them. For honeybees and paper wasps, we use #2 set screws to minimize the contact area between the push rod and the thorax. For the larger bumblebee, we use #2 hex screws. The larger mounting surface of the hex head prevented the thorax from rotating as it was compressed. Insects are mounted upside down such that their tergums are aligned with the force sensor. When mounting an insect, we use a small beads of glue to secure the dorsal and ventral surfaces of the thorax to their respective push rods. Once an insect is mounted, we lower the vertical stage until the insect’s wings begin to articulate.

We use a dynamic signal analyzer (Data Physics, ABACUS 901) both to record force and acceleration data as well as to generate excitation reference signals. The signal analyzer converts time-domain measurements to the frequency domain via a fast Fourier transform (FFT). We actuate the shaker with a “periodic chirp” from 10 - 1000 Hz over a 400 ms period with a signal width of 60%. Displacement, as estimated by twice integrating the acceleration measurement, varies with excitation frequency but is typically restricted to less than ± 1 μm to ensure that the thorax is excited in the linear range. Data is recorded at a rate of 2560 Hz, resulting in 400 FFT lines with a frequency resolution of 2.5 Hz. We utilize a Hanning window during data collection and data is averaged over at least 50 periods to reduce noise. Through this procedure, we determine the FFT *G*(*jω*) relating acceleration and force as measured at the insect tergum. We then identify the thorax’s fundamental frequency by fitting *G*(*jω*) via MATLAB’s modal fitting tool with peak picking (PP) algorithm. In instances where the peak peaking algorithm failed to fit the FRF, we opted for a least-squares rational function (LSRF) fitting algorithm. In both cases a 10-sample moving mean filter was applied to the frequency domain data set to improve the accuracy of the fit.

Once data is collected for a level of thorax compression, we compress the thorax further using the fine linear stage and repeat the experiment. For the purposes of this work, we consider a compression range of 50 - 300 *μm* in 50 *μm* increments. We found that a minimal level of compression was required to get clean FRF measurements, and that approximately 300 *μm* of dorsoventral compression was required to fully articulate the insect’s wings for the species considered. However, this maximum compression is likely higher than the thorax would experience in nature. Though we were unable to find thorax displacement measurements for the insects studied, we look to other species for an estimate. The valley carpenter bee *Xylocopa varipuncta* thorax oscillates approximately between ± 150 *μm* during flight [25]. Displacements of the stingless bee *Melipona seminigra* thorax were estimated at 20 *μm* peak-to-peak during tethered flight and up to 50 *μm* peak-to-peak during “annoyance buzzing” behavior [26]. One potential issue with using these estimates to inform our study, however, is that thorax velocity (from which displacement was estimated via integration) was measured at a single point on the thorax’s dorsal surface with laser vibrometry. Ando and Kanzaki measured the dorsal surface of a hawkmoth *Agrius convolvuli* thorax using a high-speed profilometer found displacements varied significantly along the medial-lateral direction [27]. Thus, it is likely that thorax displacements estimated via single point laser vibrometry are sensitive to the precise location of the measurement. Further, since the push rods deform the thorax in a manner dissimilar to the flight muscles, it is possible the thorax must be compressed further than typical to cause maximum wing articulation. We maintain the 50 - 300 *μm* range to ensure we capture relevant data both inside and potentially outside the anatomical range, with the understanding that induced deformations will deviate from natural deformations. We do not consider tensile loading in this study.

### 2.2. Insect Collection and Preparation

Honeybees and bumblebees were collected while foraging at Montana State University’s (MSU) pollinator garden. Paper wasps were collected during a hive removal near Bozeman, MT. All subjects were collected and transported to the laboratory immediately before experimentation and were housed in containers up to several hours. Insects were sacrificed in a sealed jar containing plaster saturated in ethyl acetate. We carefully removed legs from the ventral surface of the thorax in order to expose a smooth mounting surface for the push rods. The insect’s wings were left intact. Following euthanization, experiments lasted maximally one hour (typically around twenty minutes) to minimize dessication or other effects that may potentially affect the dynamic properties of the thorax.

## 3. Results

In this section, we use the experimental setup to investigate the linear dynamics of the insect thorax. We first compare measured thorax fundamental frequencies to reported values of wingbeat frequency. We then address the sensitivity of fundamental frequency with respect to compression. Finally, we conduct a static force-displacement test to identify the stiffness of the honeybee thorax.

### 3.1. Flapping at the Thorax Fundamental Frequency

First, we look to answer the question: do insect’s flap at the fundamental frequency of their thorax? Our results suggest that they do. At 50 μm of compression, the average thorax fundamental frequency is 368 Hz for the honeybee, 402 Hz for the paper wasp and 363 Hz for the bumblebee (Tab. 1). Previous literature indicates the wingbeat frequency is 220 - 250 Hz for the honeybee [28] and around 120 - 190 Hz for some types of bumblebees [25]. We were unable to find reported wingbeat frequencies for the paper wasp, however wingbeat frequencies for the yellowjacket (a member of the same subfamily) are 140 - 160 Hz [29]. While these measured fundamental frequencies are somewhat higher than reported wingbeat frequencies, we believe the discrepancy is largely due to artificial experimental boundary conditions. If these boundary conditions were removed, the thorax fundamental frequency likely does coincide with the insect’s wingbeat frequency.

**Table 1:**
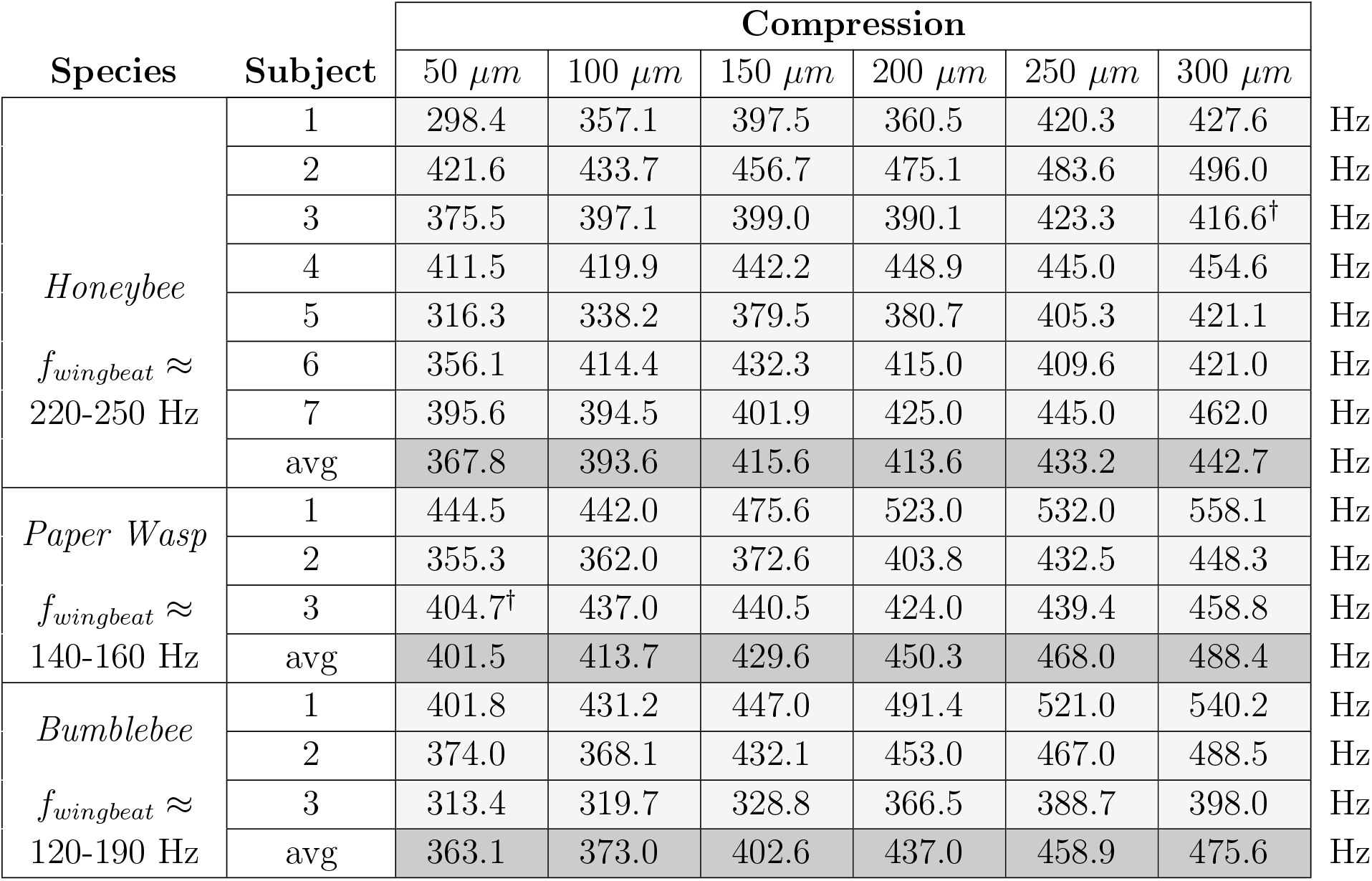
Tabulated FRF data for all insects (n=7 honeybees, n=3 bumble bees, n=3 paper wasps). The values in the gray colored boxes indicate thorax fundamental frequencies (in Hz) for a subject and compression level. † indicates that the FRF was fit via a LSRF fitting function instead of a PP fitting function. We found that some thoraxes rotated during dynamic testing, thereby changing thorax alignment and distorting FRF data. These trials were discarded.

During flight the thorax is free to oscillate, whereas in the experiment the dorsal and ventral surfaces are constrained by push rods (Fig. 2). Constraints imposed on a structure may increase its natural frequencies. To approximate the effect of boundary conditions, we develop a simple finite element (FE) model of a honeybee thorax idealized as a circular hoop and calculate numerically its natural frequencies with and without constraints (Appendix A). The FE model is not intended to replicate the thorax, but provides an approximation of how boundary conditions affect its fundamental frequency. Comparing a free hoop to a hoop with boundary conditions similar to the experiment, we find that constraints can increase the first natural frequency of the hoop by 64%. Thus, we believe the boundary conditions imposed by the experiment stiffen the insect thorax and increase its fundamental frequency non-trivially.

Constraints also help explain unanticipated trends in the data. For example, the paper wasp thorax fundamental frequency is higher than the honeybees, whereas its wingbeat frequency is lower. This may be due to the relative size of the honeybee and paper wasp thoraxes. The average thorax diameter is approximately 4.05 mm (± 0.13 mm, n=10) for the honeybee and 3.67 mm (± 0.29 mm, n=10) for the paper wasp. This implies that a greater portion of the paper wasp thorax is constrained relative to the honeybee thorax, and thus the constraint affects the paper wasp more significantly than the honeybee. Similarly, the bumblebee thorax has a fundamental frequency close to that of the honeybee despite having a lower reported wingbeat frequency. Recall that #2 hex head screws were used to compress the bumblebee whereas #2 set screws were used to compress the honeybee. The former provides a larger clamping surface, which was necessary to prevent the bumblebee from rotating during compression. The increased constrained area will dramatically increase the fundamental frequency of the bumblebee thorax.

### 3.2. Sensitivity of Fundamental Frequency to Compression

Motivated by the notion that insects are able to change wingbeat frequency, we now explore the effect of compression on thorax fundamental frequency. Interestingly, the thorax fundamental frequency tends to increase with compression for all insects considered (Tab. 1). The average fundamental frequency as a function of compression level for the honeybee is shown in Fig. 4. With the exception of the modest drop in average fundamental frequency for the honeybee between 150 *μm* and 200 *μm*, this trend occurs monotonically. This implies that the thorax’s fundamental frequency is sensitive to its instantaneous shape, which may provide a simple mechanism for the insect to adjust wingbeat frequency during flight. Prior research suggests insects from the order Diptera modulate their wingbeat frequency by tensioning their thorax via pleurosternal steering muscles [12]. While our experiment does not emulate this tensioning mechanism, it does does support the idea that thorax modifications can alter its dynamic response.

**Figure 4:**
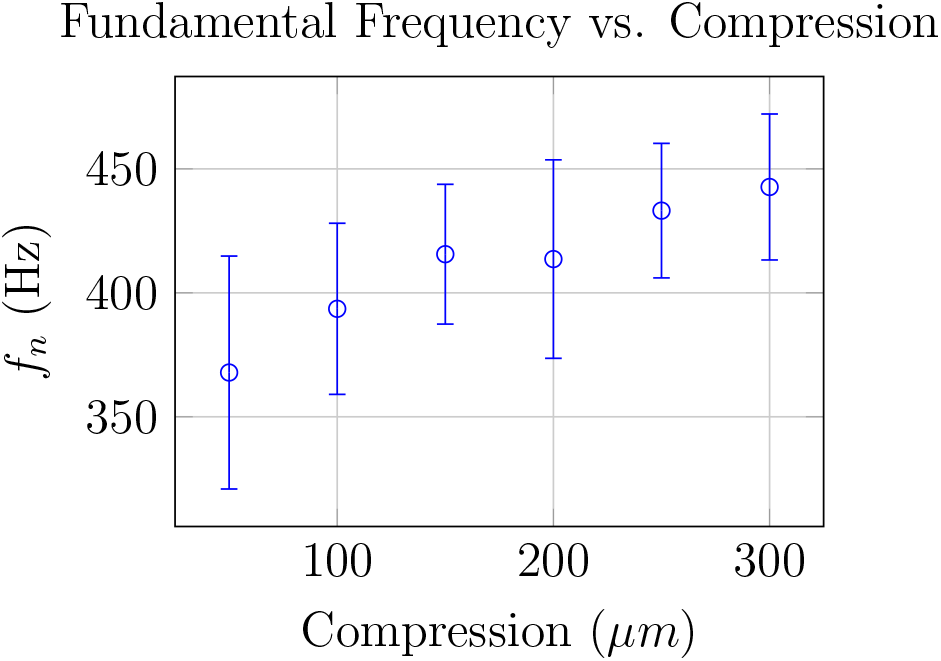
Average fundamental frequency, *f_n_*, as a function of dorsal-ventral compression for honeybee specimens (n=7). Bars show standard deviation.

The ability to modulate the thorax’s fundamental frequency is important for flight as well as other behaviors. Some Hymenoptera, including the bumblebee, perform a behavior called buzz pollination [30]. During buzz pollination, the bee will land on a flower, disengage its wings and excite its flight muscles at high frequency. The high frequency vibrations help dislodge the flower’s pollen so that the bee can collect it more easily. Flight muscle oscillations during buzz pollination occur a much higher frequency than during flight, which implies the fundamental frequency of the thorax is higher during the buzz pollination behavior. The increase in thorax fundamental frequency may result from the bee disengaging its wings, since disengaging the wings reduces the effective inertia of the wing-thorax system.

However, some bees vary their buzz pollination frequency depending on the physical attributes of the flower they are collecting pollen from [31]. This suggests that the bee can adjust its thorax fundamental frequency even while its wings are disengaged. Further, King et al. showed that during buzz polination, carpenter bees adjusted the mean position about which their thorax oscillates [25], which may indicate muscle tensioning. Thus, it is likely that both the reduction of inertia due to wing disengagement, as well as an increase in muscle tension, contribute to the high frequency oscillations observed during buzz pollination.

### 3.3. Hardening Spring Behavior

We observed that the thorax fundamental frequency increased with compression level for all insects tested. This suggests that the thorax behaves as a nonlinear hardening spring, where its stiffness increases with respect to its degree of deformation [32]. To verify this, we measure the dorsal-ventral force-displacement curve of honeybee thoraxes (n=10). The derivative of the curve with respect to displacement yields the local thorax stiffness.

We use the same general experimental set-up for static testing with two key differences: (1) the vibration shaker is replaced with a rigid support, and (2) the piezoelectric force sensor is replaced with a high-sensitivity foil-based load cell (Transducer Techniques, GSO-100). We use a National Instruments cDAQ to record force data. For the purposes of this work, we recorded the load immediately after the thorax was compressed to minimize the effect of hysteresis on measurements. We considered only increasing compression levels (loading) and did not record the unloading phase of the thorax. Results are shown in Fig. 5.

**Figure 5:**
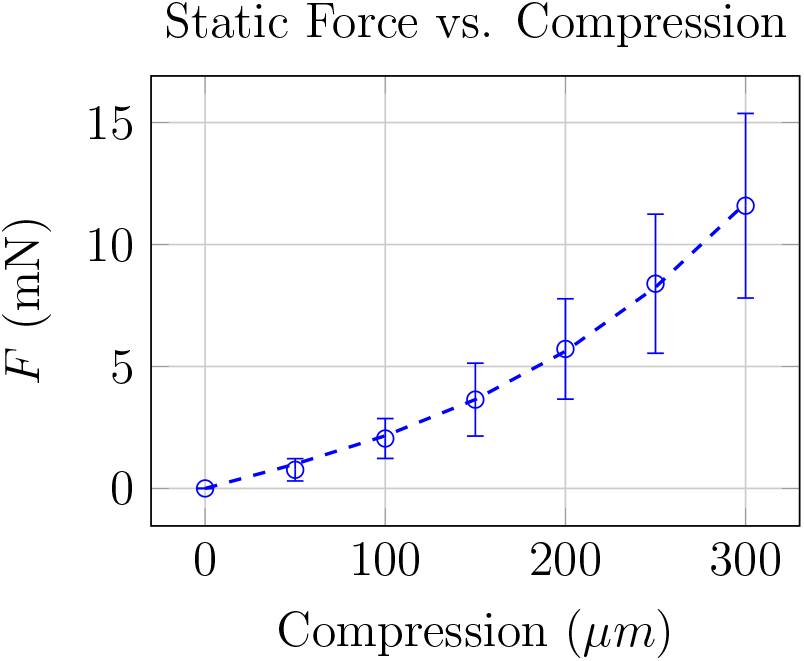
Average dorsal-ventral force-displacement curve of honeybee thorax (n=10) specimens. Bars show standard deviation.

These results verify that the honeybee thorax behaves as a hardening spring. The slope of the force-displacement, which increases with respect to thorax compression, defines the thorax’s local stiffness. We curve fit to average data from Fig. 5 to determine the thorax’s elastic force *F_sp_*(*x*) as a function of compression *x* (Fig. 3). Assuming a linear fit with an intercept of zero, the coefficient of determination R^2^ = 0.9802. For a linear-cubic fit of the form *F_sp_*(*x*) = *a*-_1_*x*^3^ + *a*_2_*x* the fit improves to R^2^ = 0.9992. We maintain the linear-cubic fit for subsequent analysis due to the higher coefficient of determination.

In general, the thorax’s fundamental frequency will be proportional to its mass and stiffness. If we assume the mass is constant, thorax’s fundamental frequency *f_n_* roughly scales with

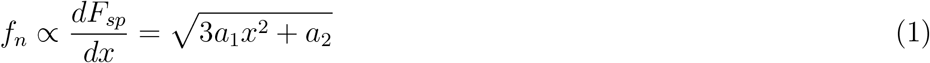

The above suggests that *f_n_* will increase linearly when x is large. To assess this, we plot the average fundamental frequency versus compression from Tab. 1 for all insects as well as a linear fit of the data in Fig. 6. In general, the linear fit models the increase in *f_n_* fairly well, with R^2^ values of 0.934, 0.994 and 0.982 for the honeybee, paper wasp and bumblebee, respectively. This finding substantiates the idea that the thorax behaves as a nonlinear hardening spring, at least about its dorsoventral axis.

**Figure 6:**
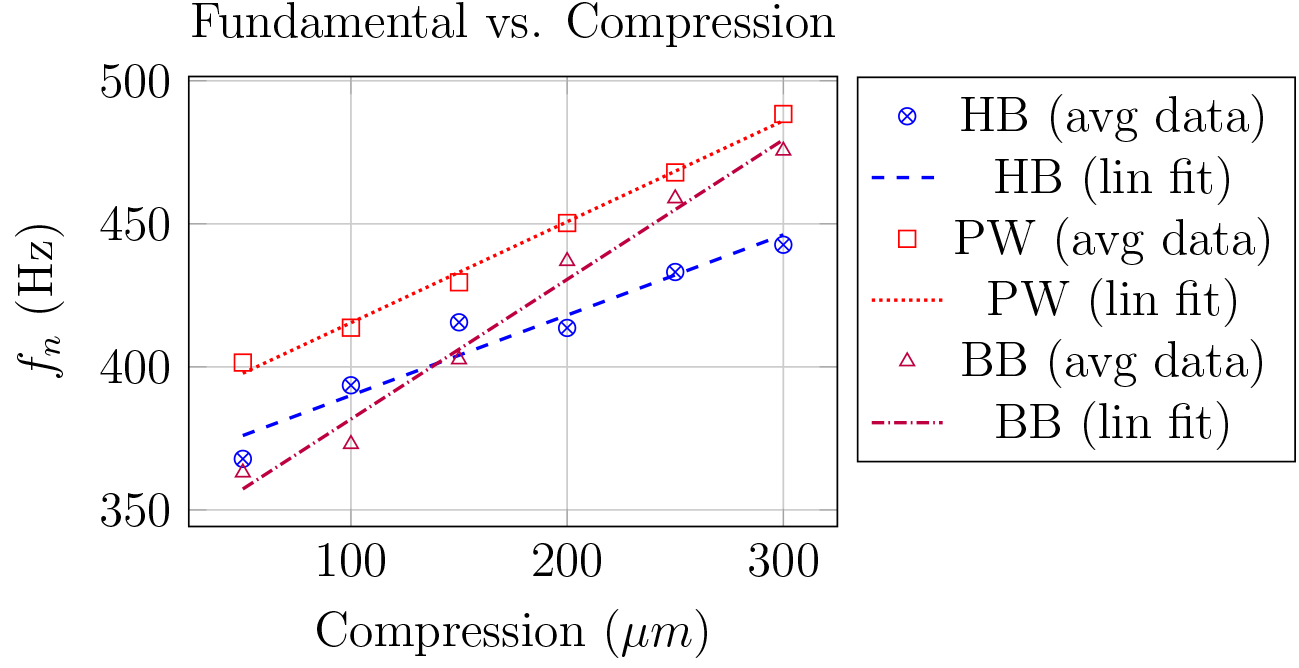
Average fundamental frequency, *f_n_* as a function of dorsal-ventral compression for all insects. HB denotes honeybee, PW denotes paper wasp, BB denotes bumblebee. Linear regression fits are shown for each data set.

## 4. Discussion

In this work, we investigated the hypothesis that many insects flap at the fundamental frequency of their thorax. We considered insects specifically of the order Hymenoptera, including the honeybee *Apis mellifera*, European paper wasp *Polistes dominula*, and bumblebee *Bombus huntii*. We characterized the linear frequency response function relating input force to output acceleration for several thoraxes. We found that, when compressed minimally, measured thorax fundamental frequencies were approximately 50-150% higher than the insects’ reported wingbeat frequency. However, the experimental assembly constrains the thorax in a manner that may significantly stiffen it, thereby increasing its fundamental frequency relative to the free boundary conditions that would be experienced during flight. If experimental boundary conditions were removed, it is plausible that the insects considered indeed flap at their thorax’s fundamental frequency. Next, we compressed the thorax and measured its FRF to identify if the thorax’s fundamental frequency was sensitive to changes in geometry. We found that the thorax’s fundamental frequency increased with respect to compression. This may be a mechanism that insects utilize to change their wingbeat frequency during flight or other behaviors. Further analysis of this research, including the energetic benefits of flapping at resonance and the potential nonlinear dynamics of the thorax, is discussed in the following sections.

### 4.1. The Energetics of Flapping at the Thorax Fundamental Frequency

Our results that flying insects likely do flap at the fundamental frequency of their thorax. Why is this beneficial? Consider the honeybee thorax FRF shown in Fig. 7. At the thorax’s fundamental frequency, the FRF magnitude |*G*(*jω*)| increases dramatically while the phase undergoes a 180° shift. The peak of |*G*(*jω*)| occurs due to a minimum in the force magnitude |*F_opp_*(*jω*)|, which implies less force is required to oscillate the thorax at this frequency. In general, the energy required to flap the wings is equivalent to the work *W* done by the thoracic force, or

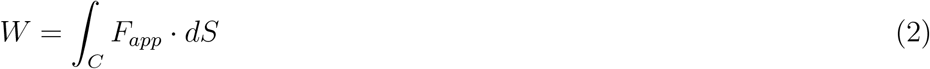

**Figure 7:**
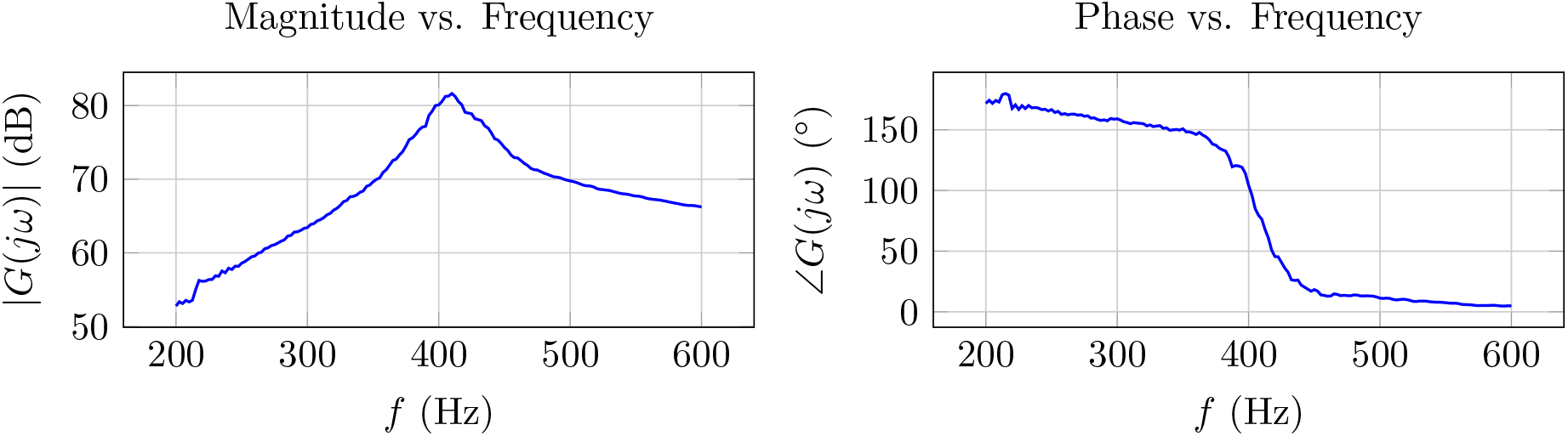
Frequency response function magnitude and phase of a honeybee thorax (subject 4) compressed 50*μm*. The sharp magnitude peak and 180° phase shift indicate a resonance at 411 Hz. Data is unfiltered.

where *C* is the trajectory of which the force acts and *dS* is the differential distance. In general, the thorax trajectory *C* is fixed because a certain amount of deformation is required to articulate the wings. Then, in order to reduce the energy required to flap the wings, the applied force must be lowered. This is achieved via flapping at resonance. In a sense, flapping in this manner maximizes the work done by conservative forces, namely the passive elastic force the thoracic exoskeleton (and potentially the passive elasticity of the flight muscle). These forces are responsible for slowing the wing down or “braking”. Without thorax elasticity, energy would have to be invested both to accelerate and decelerate the wing. Therefore, insects may flap at the fundamental frequency of their thorax in order to reduce the energetic requirements of flight.

### 4.2. Thorax as a Duffing Oscillator

The experiments conducted here considered low amplitude excitation and were designed only to force the thorax in its linear range. However, the results of this research suggest that the thorax may behave as a weakly non-linear oscillator when excited at larger amplitudes. Notably, the thorax behaves as a nonlinear hardening spring, which implies that its stiffness and fundamental frequency increase with respect to aconstant level of compression. The cubic stiffness nonlinearity is the hallmark of the Duffing oscillator [32]. With linear damping and external harmonic forcing, the equation of motion for the Duffing oscillator is

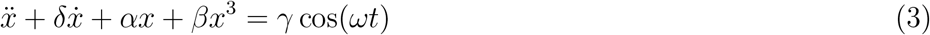

where *δ* is the linear viscous damping coefficient, *α* is the linear stiffness coefficient, *β* is the cubic stiffness coefficient, *γ* is the excitation amplitude and ω is the harmonic forcing frequency. The behavior of the Duffing oscillator is well studied and may provide insight into the nonlinear mechanics of the insect thorax.

One notable feature of the Duffing oscillator is that, when subject to a forcing with periodicity of *ω*, it responds both at *ω* and integer harmonics thereof. Hrncir et. al observed such harmonics when measuring the thorax during “annoyance buzzing” in bees [26]. Because thorax deformation also drives flapping, it is possible these harmonics affect flapping kinematics as well. Recordings of hawkmoth *Manduca sexta* wingbeat kinematics indicate the presence of higher-order harmonics of the flapping frequency in all three wing rotations [33]. Higher-order harmonics of the flapping frequency appear to play a significant role in wing deformation and the corresponding energetic and aerodynamic efficiency of flight [34—36].

A second essential feature of the Duffing oscillator is its amplitude dependence. Unlike linear oscillators, the frequency at which the Duffing oscillator exhibits the largest response depends on excitation amplitude. This diverges from linear oscillators where the fundamental frequency depends on only the properties of the structure and not the excitation amplitude. This amplitude dependence may have some interesting implications for artificial and biological flight. In fruit flies, stroke amplitude has been shown to increase with wingbeat frequency [22]. If the fruit fly thorax behaves as a Duffing oscillator, an increase in thorax deformation amplitude would increase both stroke amplitude as well as the thorax’s fundamental frequency. This would increase the range of frequencies that correspond to a resonant response.

While this is an interesting idea, it is unclear if insects leverage nonlinearity and amplitude dependence to affect thorax resonant frequencies. Unlike fruit flies, honeybees typically maintain their wingbeat frequency while adjusting their stroke amplitude to produce additional lift while climbing [37]. The increase in stroke amplitude could stem from larger thorax deformations, which would imply that the thorax resonant frequency is insensitive to increases in excitation amplitude. However, it is also possible that modifications to the wing hinge kinematics cause increased stroke amplitude while thorax deformation amplitude is maintained. This latter scenario would not preclude the thorax from behaving as a nonlinear oscillator.

Nonetheless, the amplitude dependent behavior of the Duffing oscillator may be desirable for FWMAVs to mimic. Many smaller FWMAVs rely on resonance between the coupled wing and actuator (often piezoelectric) to realize large stroke amplitudes [4]. While this is an effective way to amplify small actuator displacements, it permits only a narrow range of permissible flapping frequencies. Once the flapping frequency deviates significantly from the system’s fundamental frequency, its response amplitude and wing rotation attenuate starkly. In contrast, leveraging an amplitude-dependent nonlinear system will increase the permissible range of flapping frequencies, because the system’s fundamental frequency can be increased by strengthening actuator amplitude. This may provide an efficient mechanism to initiate energy efficient climbing flight, because both the stroke amplitude and wingbeat frequency increase simultaneously while the resonant behavior of the system is preserved.

## Acknowledgments

This manuscript was supported in part by the National Science Foundation under award No. CBET-1855383. Any opinions, findings, and conclusions or recommendations expressed in this material are those of the author(s) and do not necessarily reflect the views of the National Science Foundation. The authors declare no conflict of interest regarding the publication of this article. We would like to thank Keith Fisher for retrieving the paper wasp nest, Andrew Mountcastle for his useful conversation on buzz pollination, and Data Physics Inc. for providing the dynamic signal analyzer.

## APPENDIX A Simplified Finite Element Modeling

We develop a simplified finite element model (Fig. 8) to estimate how experimentally boundary conditions affect the fundamental frequency of the insect thorax. The simple model is not intended to reproduce experimental findings, but instead provide insight into how the push rod constraints (Fig. 2) influence the thorax’s dynamic response.

**Figure 8:**
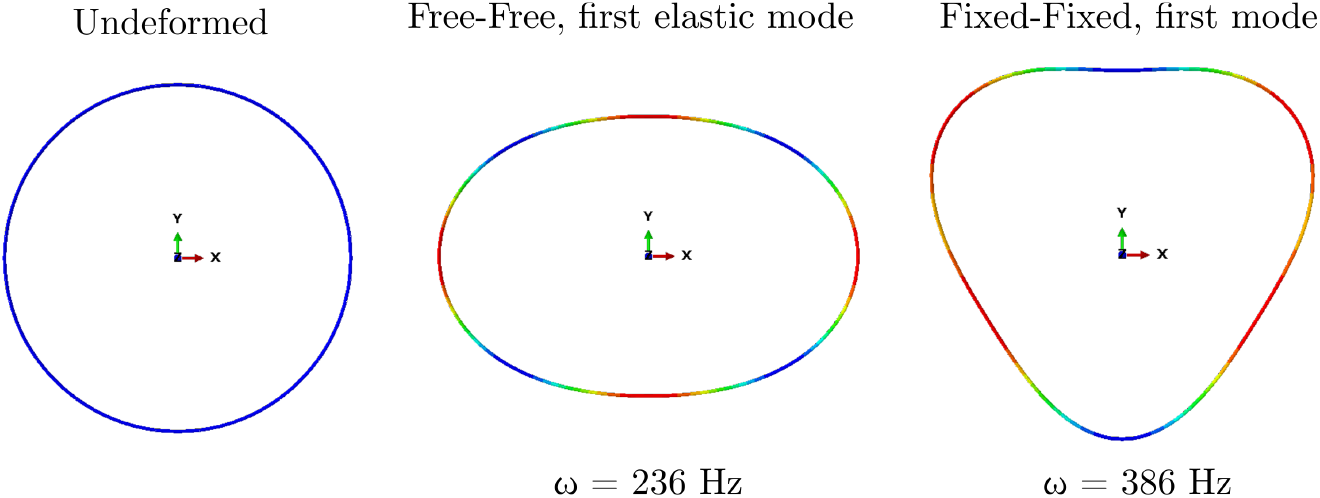
Simplified FEA model to demonstrate the effect of boundary condition on the insect thorax frequency response. The fixed-fixed model has a first natural frequency of 150 Hz higher than the free-free model.

We idealize a transverse cross section of the thorax as a planar, uniform thin-walled hoop with a radius of 3 mm. We model only half the hoop and assume symmetry about a vertical line bisecting the hoop. The hoop wall is solid and circular with a radius of 10 μm, density of *ρ* = 0.858 g/mm^3^ and Young’s modulus *E* = 8.5×10^12^ Pa. The modeled hoop section is discretized into 50 beam elements of equal lengths, which is a sufficient to show convergence of the structure’s first several natural frequencies.

We consider free-free boundary conditions, which is representative of free flight, and fixed-fixed boundary conditions, which is similar to those imposed during the experiment. For the free-free case, single points at the top and bottom of the circle are constrained such that they can only translate in the *y* direction; *z* axis rotation and *x* translation are restricted. These are necessary boundary conditions when assuming the thorax is symmetric about *y* axis. For the fixed-fixed case, the points at the top and bottom of the circle are constrained to have no rotation or translation. For both cases, we conduct numerical modal analysis to identify the first two natural frequencies and corresponding mode shapes of the idealized thorax (Fig. 8). Note that the first natural frequency of the free-free model will be zero, which indicates rigid body translation of the structure in the *y* direction. For this reason, we compare the first non-zero natural frequency for both boundary condition cases.

Indeed, the fixed-fixed thorax has a first non-zero natural frequency of 386 Hz, which is 150 Hz higher than the free-free thorax natural frequency of 236 Hz. This strongly suggests that the experimental boundary conditions may significantly increase the measured the thorax natural frequency. Increasing the clamped area tends to increase the natural frequencies even further. Moreover, the vibration mode shape itself is altered. Whereas the first vibration mode of the free-free thorax is symmetric about the *x* axis, this symmetry is broken for the fixed-fixed case. Given these limitations of the experimental set-up, future efforts will be devoted to repeating portions of this experiment with boundary conditions more representative of free flight. Nonetheless, the data collected here supports the notion that many insects with asynchronous flight muscles flap at the fundamental frequency of their thorax.

## Notes

#### Summary of Updates

We have revised the spelling "Hymonoptera" to the correct spelling "Hymenoptera"

